# The Evolution of Heteroresistance via Small Colony Variants in *Escherichia coli* Following Long Term Exposure to Bacteriostatic Antibiotics

**DOI:** 10.1101/2023.10.30.564761

**Authors:** Teresa Gil-Gil, Brandon A. Berryhill, Joshua A. Manuel, Andrew P. Smith, Ingrid C. McCall, Fernando Baquero, Bruce R. Levin

## Abstract

Traditionally, bacteriostatic antibiotics are agents able to arrest bacterial growth. Despite being traditionally viewed as unable to kill bacterial cells, when they are used clinically the outcome of these drugs is frequently as effective as when a bactericidal drug is used. We explore the dynamics of *Escherichia coli* after exposure to two ribosome-targeting bacteriostatic antibiotics, chloramphenicol and azithromycin, for thirty days. The results of our experiments provide evidence that bacteria exposed to these drugs replicate, evolve, and generate a sub-population of small colony variants (SCVs) which are resistant to multiple drugs. These SCVs contribute to the evolution of heteroresistance and rapidly revert to a susceptible state once the antibiotic is removed. Stated another way, exposure to bacteriostatic drugs selects for the evolution of heteroresistance in populations previously lacking this trait. More generally, our results question the definition of bacteriostasis as populations exposed to bacteriostatic drugs are replicating despite the lack of net growth.

## Introduction

Antibiotics can be broadly classified as being bactericidal or bacteriostatic based on whether they kill bacteria or simply arrest their growth ^1^. However, it is known that at high drug concentrations bacteriostatic drugs can and do kill bacteria. Intuitively, it would make sense to treat an infection with drugs that readily kill the infecting bacteria, the bactericidal drugs, and thereby eliminate the reliance on the host’s immune system to clear the infection, as would be the case with bacteriostatic antibiotics. For this reason, bacteriostatic drugs have been considered “weaker” than bactericidal drugs and are not recommended for the treatment of severe infections or infections in immunodeficient patients ^2,3^.

This distinction between bactericidal and bacteriostatic drugs is manifest in quantitative experimental studies of the pharmacodynamics (PD) of antibiotics and bacteria. These studies focus on the rates and dynamics of the drug’s ability to kill exposed populations of bacteria ^4-7^. Many of the studies concerning why antibiotics fail to control bacterial infections have focused on phenomena solely belonging to bactericidal antibiotics such as persistence and tolerance, that is the ability to survive exposure to high concentrations of an antibiotic of a subpopulation or the whole bacterial population, respectively ^6,8-10^. These traits are qualities only associated with bactericidal drugs: the bacteriostatic drugs do not kill, and consequently the minority surviving or slower-dying populations are not revealed. With much of the clinical application of antibiotics focusing on bactericidal drugs and the majority of the research on the PD of antibiotics and bacteria also focusing on bactericidal drugs, PD research on bacteriostatic drugs has been relatively neglected.

However, in recent years, clinicians have given less importance to the antibiotic’s ability to kill bacteria *in vitro* and instead have focused on the outcome of treatment with these drugs. By this criterion, in many cases bacteriostatic antibiotics are as effective as bactericidal even in severe infections, with the possible exception of immunosuppressed patients ^2,11^. The increase in use of bacteriostatic antibiotics could reduce the selection pressure for resistance to bactericidal agents used in these critical infections. However, the shift to using bacteriostatic antibiotics requires the development of quantitative measures of the PD of antibiotics that arrest the growth of, rather than kill, bacteria ^12^. One cannot solely characterize the PD of bacteriostatic antibiotics by the minimum concentration required to prevent the replication of exposed bacteria, the MIC ^13^. Furthermore, the evolution of genomic resistance for these agents is rare and the resistant traits tolerance or persistence that occur in bactericidal antibiotics are difficult to define and perhaps impossible to detect with these bacteriostatic agents ^14,15^. This raises questions about the population and evolutionary dynamics of bacteria confronted with these drugs. If the bacteria exposed to bacteriostatic antibiotics are not replicating, one would not expect them to evolve. Therefore, more considerations of the pharmaco-, population, and evolutionary dynamics of bacteriostatic antibiotics are needed both academically and clinically.

In this study, we explore the pharmaco-, population, and evolutionary dynamics of *Escherichia coli* exposed to two bacteriostatic antibiotics of different classes, chloramphenicol (CHL) and azithromycin (AZM), over 30 days. While resistance did evolve over the 30 days, we did not detect any previously described resistance mechanisms, which include antibiotic efflux, decrease in membrane permeability, or antibiotic inactivation for CHL ^16^; nor the overproduction of efflux pumps or mutations in genes encoding the 23S rRNA subunit for AZM ^17^. The results of our experiments provide evidence that: (i) long-term exposure to these ribosome-targeting bacteriostatic antibiotics does not change the net density of exposed populations, (ii) despite the fact that the population’s net density does not change, bacteria exposed to these drugs replicate, evolve, and generate small colony variants, and (iii) the selective pressure mediated by these drugs favors the evolution of heteroresistant populations, i.e. the emergence of resistant minority populations, in a strain previously lacking this trait.

## Results

### Long Term Exposure to Bacteriostatic Antibiotics

We begin our investigation into the effects of long-term exposure to ribosome-targeting bacteriostatic drugs by evaluating the impact that these agents have on bacterial survival over 30 days. We exposed four independent cultures of ∼10^5^ CFU/mL of *E. coli* MG1655 in glucose-limited minimal media to super-MIC (Minimum Inhibitory Concentration) concentrations of CHL and AZM for 30 days without transferring or diluting (Figure 1). One thousand μg/mL of glucose was used since this concentration allows cultures to grow to maximum stationary densities while avoiding nutrient limitation. The MIC of CHL and AZM with MG1655 were estimated by broth microdution in this glucose-limited minimal media and found to be 6.25 ug/mL for both drugs ^13^. Super-MIC concentrations of each drug that were shown to be bacteriostatic with minimal killing and/or growth were used in the experiment (Supplementary Figure 1). Cultures exposed to these drugs were sampled every 5 days. We observed that over the course of the experiment, the control cultures containing no drugs reached their maximum stationary phase density of approximately 10^9^ CFU/mL and then went down by approximately two logs over the course of 30 days, while the densities in the cultures containing the drugs remained stable, with at most a half-log change in density in the drug-treated populations. Notably, in drug-treated cultures where the bacteria were not lost, two distinct colony morphologies emerged. The colonies were either similar to the ancestral wild-type *E. coli* or were much smaller bacterial colonies, small colony variants (SCVs). This evolution occurred while under strong selective pressure from these drugs. There was no change in colony size in the drug-free controls, with a limit of detection of 10^−3^ SCVs per colony of the ancestral population. These SCVs started to emerge at day 10 of the long-term experiment in cultures exposed to CHL and at day 15 in cultures exposed to AZM. Their frequency increased from 2-5% when they were first observed in the presence of both antibiotics to 10-44% in CHL and 10-50% in AZM by day 30. To assess the maintenance of the activity of the antibiotics after 30 days, bacteria resistant to the antibiotic in each culture was added at approximately 10^6^ CFU/mL and over the course of 24 hours each culture grew approximately three orders of magnitude; while when the susceptible, wild-type *E. coli* strain was added to these 30 days supernatants it was not able to increase in density (Supplementary Figure 2). This residual growth indicates that at 30 days the antibiotic is still at a super-MIC concentration and thereby is the limiting factor for growth in the long term experiments. Moreover, 0- and 30-day supernatants were filtered, and the amount of antibiotic present tested through a disk diffusion assay. The diameter of both time points was 1 cm for the CHL and 2.8 cm for the AZM supernatants confirming the stability of both antibiotics during the long term experiments.

**Fig 1.**
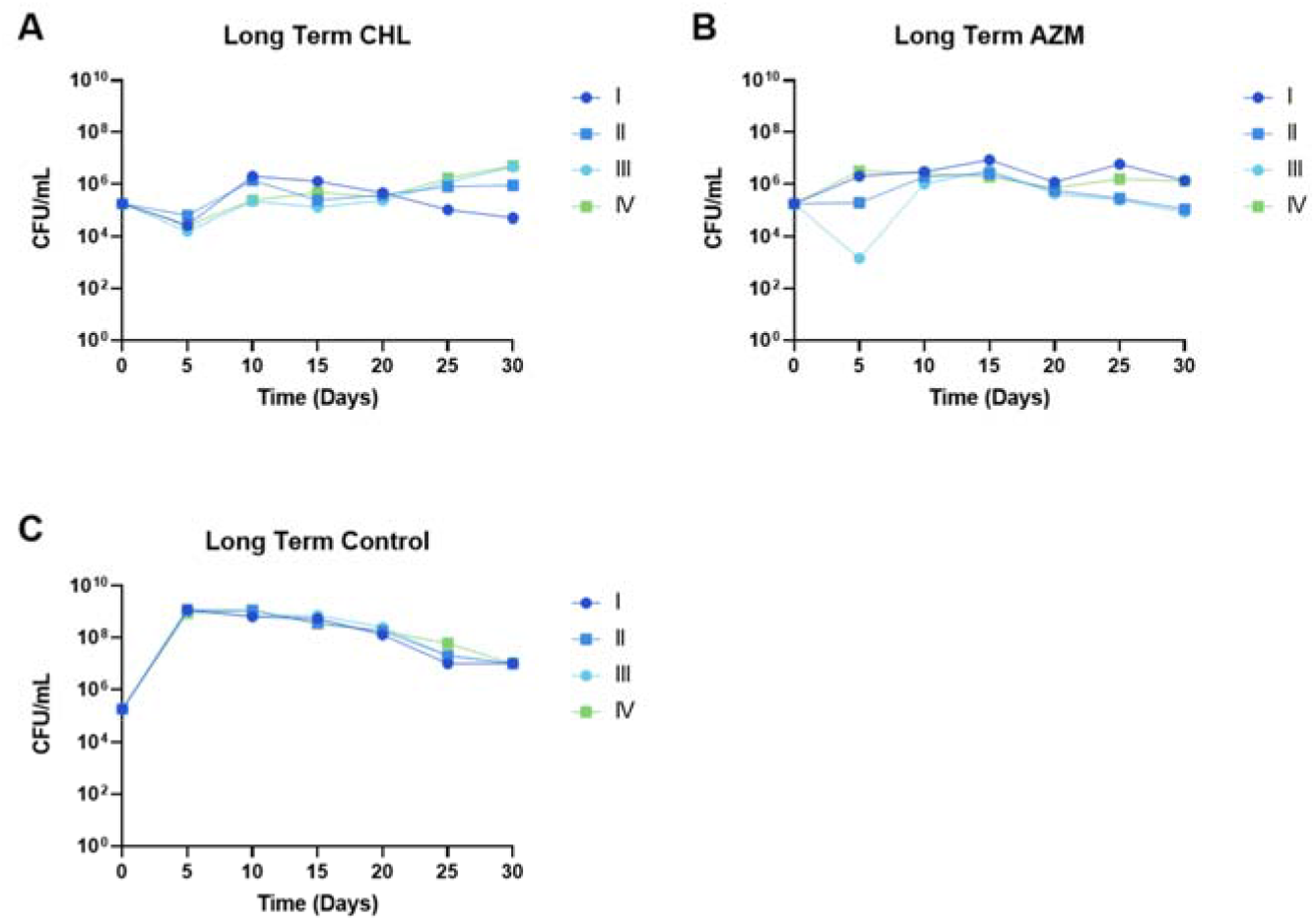
Long term exposure of *E. coli* to bacteriostatic drugs. Density in CFU/mL of *E. coli* MG1655 measured every 5 days for 30 days of 4 independent biological replicas (I-IV). **(A)** *E. coli* exposed to 4x MIC CHL (chloramphenicol); **(B)** *E. coli* exposed to 3x MIC AZM (azithromycin); **(C)** Drug-free control.

To determine how evolution is occurring in the apparent absence of net growth in the presence of the antibiotic, we performed a long term experiment using a conditionally non-replicative plasmid ^18^ to identify if growth is occurring at the same rate as death (Figure 2). After 10 days the plasmid frequency decreased by 100-fold, and after 20 days the plasmid frequency was near the limit of detection. That means that the plasmid containing cells are progressively diluted (as each cell division gives rise to a plasmid-free descendant) and indicates that the population is growing, replicating at least once a day and dying at the same rate. To explain how these results are consistent with the hypothesis that is occurring at the same rate as death, we constructed a mathematical and computer-simulation model of the plasmid being lost both with and without bacteriostatic drugs (Supplementary Text, Supplementary Equations 1-4, and Supplementary Figure 3).

**Fig 2.**
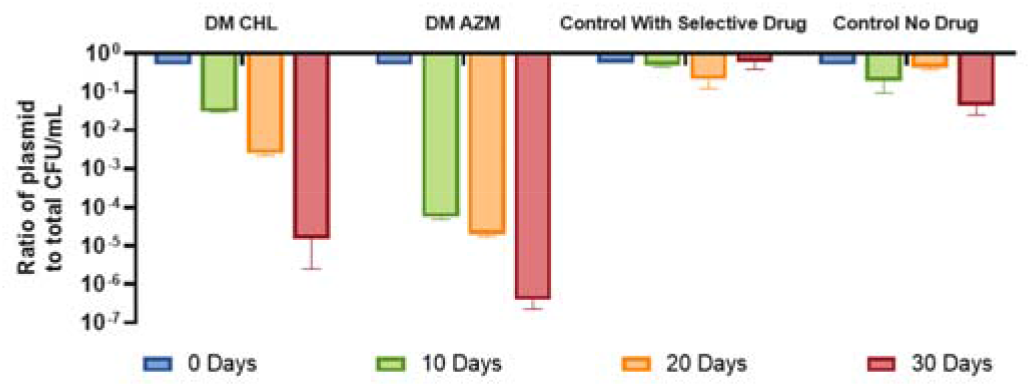
Long term experiment with a non-replicative plasmid. Ratio of the plasmid-containing cells to the total number of cells in CFU/mL of *E. coli* MG1655 with the non-replicative plasmid pAM34 which bears an ampicillin resistance cassette. The total cell density and density of cells bearing the plasmid were measured every 10 days for 30 days of 4 independent biological replicas for four conditions: i) minimal media with CHL (chloramphenicol), ii) minimal media with AZM (azithromycin), iii) minimal media with ampicillin– maximal plasmid retention, iv) minimal media with no antibiotic– maximal plasmid loss. In the absence of CHL or AZM the initial cultures can only grow two logs until arriving to a maximum stationary phase, explaining the maximum loss of two log in the absence of ampicillin.

### Small Colony Variants Characterization

To determine what these SCVs are, we isolated 6 independently generated SCVs of MG1655 (CHL A, CHL B, CHL C, AZM A, AZM B, and AZM C) and characterized them phenotypically and genotypically. Firstly, to determine if the SCVs are a form of resistance that has emerged over the long-term experiment, their MIC to the drugs they were previously exposed to (Figure 3). Each SCV showed at least an 8-fold change over the ancestral MG1655’s MIC. On the other hand, normal-sized colonies from each culture were isolated and changes in MIC respecting the ancestral MG1655 were not observed in any of them. Each SCV has a distinct antibiotic susceptibility profile in terms of collateral susceptibilities and cross-resistances (Supplementary Figure 4).

**Fig 3.**
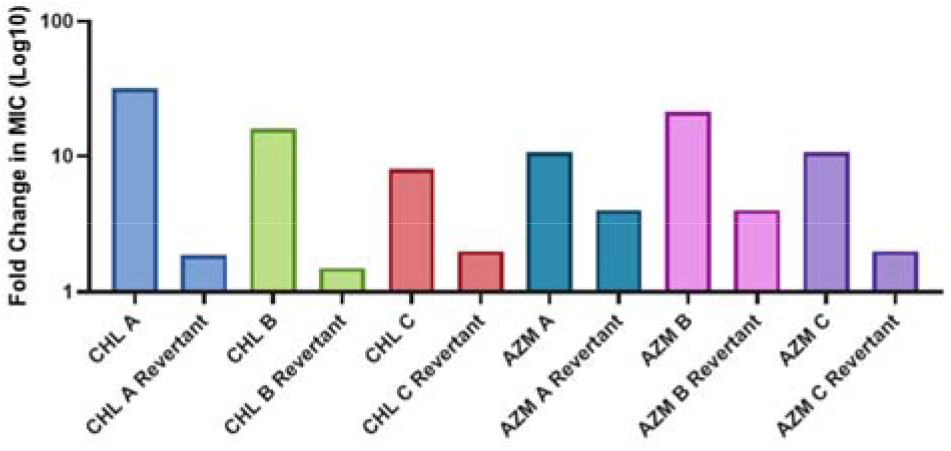
MIC of the SCVs to their respective drugs. Three SCVs were isolated from each condition from day 30 of the long term experiment, grown up in 1.5x *E. coli* MG1655’s MIC for these respective drug, and then E-tested. Fold change respecting the MIC of the final evolved control populations, whose MIC was 6.25 μg/mL as it was at the start of the long-term experiment.

Notably, although these SCVs are resistant to the bacteriostatic antibiotic to which they were exposed, there is very little net growth in the long term culture over the course of 30 days. To determine why a marked increase in density does not occur despite the evolution of resistance, we performed OD-based growth experiments of the SCVs with different concentrations of antibiotics. We found that even though these mutants are resistant, their growth rates and the maximum optical densities decreased proportionally to the drug concentration and their lag time was substantially increased (Supplementary Figure 5). This result is consistent with previous observations ^13^.

Antibiotic resistance in the small colonies obtained from both CHL and AZM cultures was unstable, that is, when streaked on LB plates without the drug the MIC of the SCVs decrease. After the genotypic characterization of these SCVs we do find genetic differences in most of them (Supplementary Table 1), but we could not find a clear mechanism that would explain this resistance. While we could not identify causal genetic differences between the SCVs and their ancestor MG1655, since both CHL and AZM prevent translation, we sought to determine if there were differences in the rate or total level of translation. Testing the level of a constitutively expressed enzyme, beta-galactosidase, we found that the six SCVs had higher levels of translation than the background MG1655 (Figure 4). Moreover, we found that the rate of translation decreases in a concentration-dependent manner to both CHL and AZM (Supplementary Figure 6). This is consistent with recent observations indicating partially compromised translation as a result of exposure to tetracycline, a translational inhibitor, which potentially accounts for the collateral susceptibility to this drug observed in Supplementary Figure 4 ^19^. Therefore, at the drug concentrations used in the long-term experiments, these SCVs are readily selected for over the ancestor, not only for their level of antibiotic resistance but also for their level of translation, explaining their ability to increase in frequency when rare.

**Figure 4.**
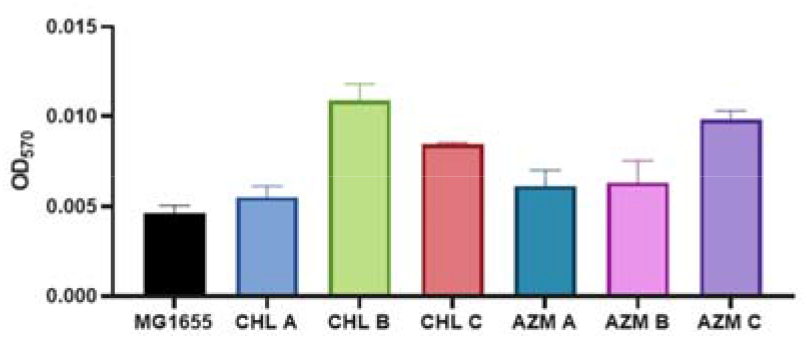
An assay of beta-galactosidase as a proxy for translation level. The beta-galactosidase levels as determined by absorbance for the ancestral *E. coli* MG1655 and the six SCVs. Shown are the means and standard deviations of three independent biological replicates.

### Heteroresistance

Antibiotic heteroresistance (HR) is defined as, “a phenotype in which a bacterial isolate contains subpopulations of cells that show a substantial reduction in antibiotic susceptibility compared with the main population”, and is detected via a population analysis profile (PAP) test ^20^. The revertant populations obtained from these SCVs have a lower MIC than those of their small colony ancestor. These revertant populations are capable of rapidly regenerating the SCVs, which have a higher MIC – meeting the definition of antibiotic HR. The ancestral *E. coli* MG1655 is not capable of generating resistant subpopulations, as shown via PAP test (Supplementary Figure 7 Panels A and B). In Figure 5, we show PAP tests of CHL B (Panel A) and AZM C (Panel B). Both SCVs are shown to be heteroresistant based on the above criteria. Moreover, highly resistant colonies isolated from these PAP tests were found to be unstable and revert to the initial SCV state in 15 days, as shown by a decrease in the MIC (Fig. 3) as well as a decrease in the frequency of antibiotic-resistant subpopulations (Fig. 5). As shown in Supplementary Figure 7 Panels C and D, all SCVs obtained from the long term experiment meet the criteria for HR. Pursuing the genetic basis of this reversion we performed a genotypic characterization of these revertants. We did find extra mutations in CHL B and AZM B (Supplementary Table 2), but once again we did not find mutations that can explain reversion.

**Fig 5.**
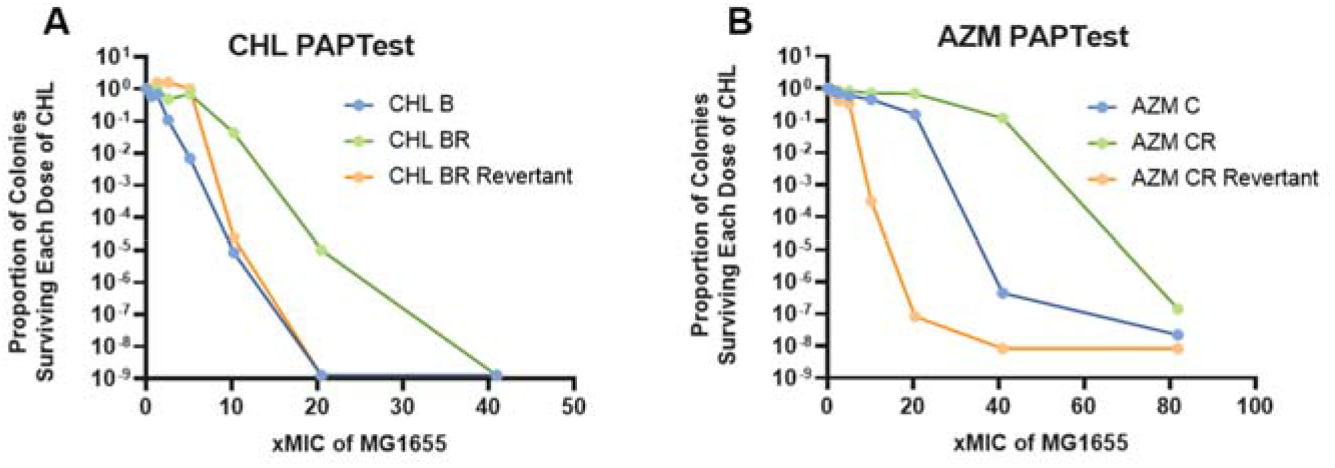
PAP tests of a CHL and an AZM SCV. **(A)** PAP tests of a chloramphenicol SCV (blue line), the most resistant isolate of this clone (green line), and the most resistant isolate after being grown without antibiotic pressure for 15 days (orange line). **(B)** PAP tests of an azithromycin SCV (blue line), the most resistant isolate of this clone (green line), and the most resistant isolate after being grown without antibiotic pressure for 15 days (orange line).

### Mathematical Model and Computer Simulations

To explore the dynamics of the emergence of the SCVs and subsequently the evolution of HR described in the above experimental results, we constructed a mathematical and computer-simulation model. In particular, we are interested in explaining the timeframe and conditions in which the SCVs arise, why the SCVs fail to ascend to a nutrient-limited density, and why the evolution of HR is stable. In Figure 6 we depict the model graphically and in the Supplementary Text and Supplementary Equations 5-11 we describe the model and its equations.

**Fig 6.**
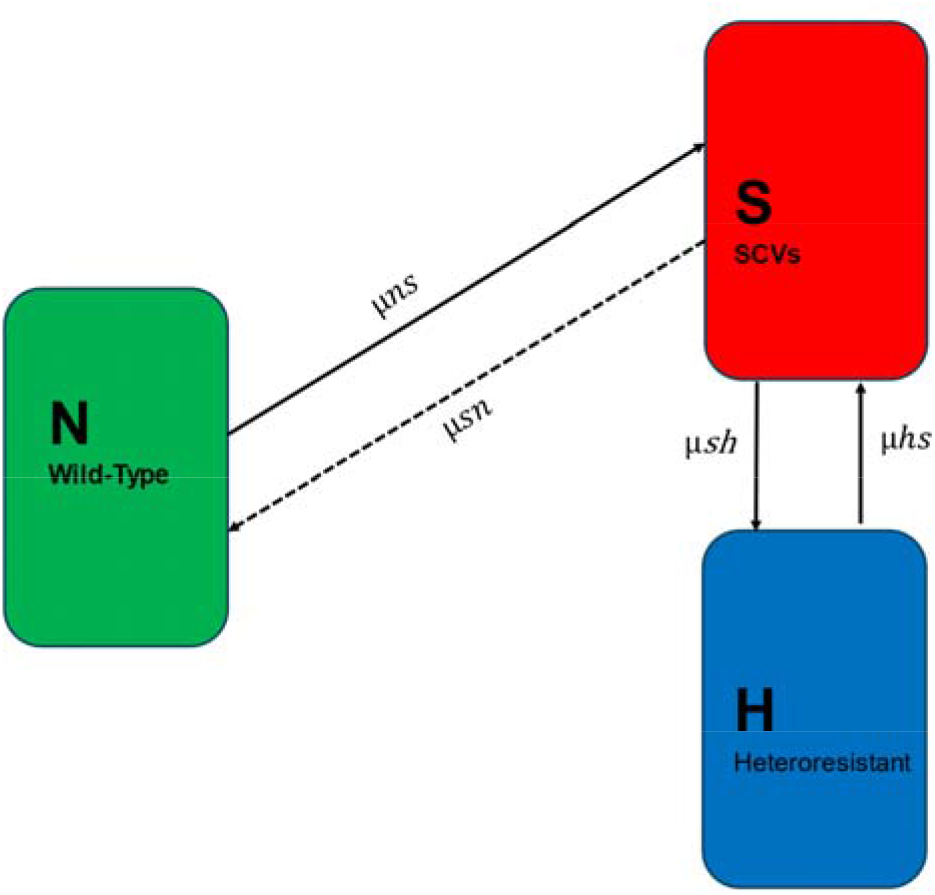
Diagram of a semi-stochastic model of the evolution of HR. The variables N, S, and H are, respectively, the wildtype *E. coli* MG1655, SCVs, and the heteroresistant bacteria in cells/mL. The parameters μ_ns_, μ_sn_, μ_sh_ and μ_hs_ are the transition rates, per cell per hour, between the different states.

In Figure 7A we illustrate how the presence of an antibiotic selects for the emergence and ascent of resistant SCVs and a heteroresistant population from an initially wild-type population. In Figure 7B, we show that an initial population of SCVs when grown without antibiotics will rapidly transition and give rise to a heteroresistant population which ascends and becomes limited by the resource. In Figure 7C, we show that exposing the heteroresistant population to the drug more rapidly selects for the emergence and dominance of reistant SCVs than when exposing the wild type population to the same concentraton of the drug. In Figure 7D, we show the changes in average MIC for the three scenarios depicted in panels A, B, and C. In Supplementary Figure 8 we show the predicted dynamics of the above model with differing concentrations of the treating antibiotic. Notably, the emergence of the SCVs and HR is dependent on this concentration—critically, the drug concentration cannot exceed the MIC of the most resistant population. For a more detailed consideration of this result and the impact that having multiple resistant states has on the emergence of HR, see ^21^. Additionally, in Supplementary Figure 9, we explore the sensitivity of our model to the governing parameters: the transitions rates and relative fitness costs. Variations in the transition rates from the wild-type to the SCVs (Supplementary Figure 9A and B) simply alter the rate at which SCVs emerge and dominate, without changing the overall outcome. Similarly, adjusting the transition rates from SCVs to the heteroresistant population (Supplementary Figure 9C and D) only affects the frequency and timing of SCV dominance. Lastly, differing fitness costs of the SCVs (Supplementary Figure 9E and F) merely impact the time required for SCV dominance. Together, these mathematical simulations illustrate that our observations hold true across a wide range of biologically meaningful parameter values.

**Figure 7.**
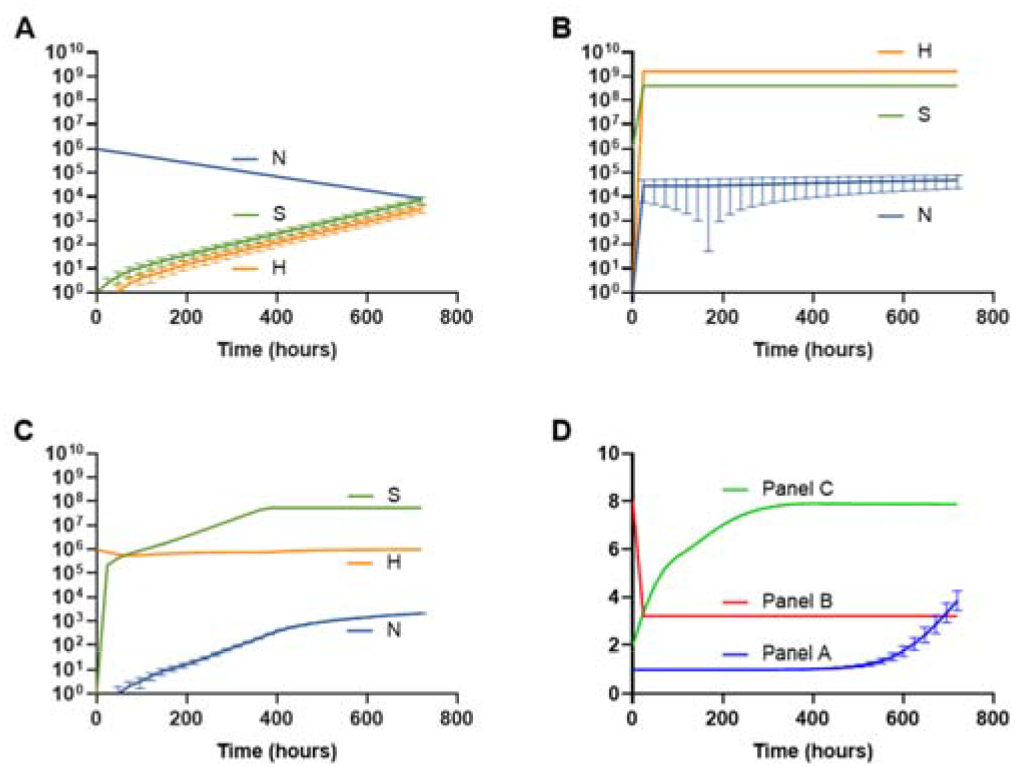
Computer simulations of the proposed model of evolved HR. Parameters used for these simulations are *e*= 5x10^−7^ μg/cell; *v*_*maxN*_ = 1.0, *v*_*maxS*_ = 0.5, *v*_*maxH*_ = 1.0 per cell per hour; *v*_*min*_ = -0.01 per cell per hour; K=1; k=1; MIC_N_=1.0, MIC_S_=8.0, MIC_H_=2.0 μg/ml; *da*= 0 μg/hour; μ_ns_=1x10^−8^, μ_sn_=1x10^−8^, μ_sh_=1x10^−3^, μ_hs_= 1x10^−3^ per cell per hour. The variables N, S, and H are, respectively, the wildtype *E. coli* MG1655, SCVs, and the heteroresistant bacteria in cells/mL. Note, points which do not appear to have error bars have standard deviations which are too small to be graphed. One hundred independent runs of the models were carried out in each condition. **(A)** H and S selection from an initial sensitive population. Parameters are N=1x10^6^ cells/mL; A=3 μg/mL. **(B)** H selection from an initial SCV population in the absence of the drug. Parameters are S=1x10^6^ cells/mL; A=0 μg/mL. **(C)** S selection from an initial H population in the presence of the drug. H=1x10^6^ cells/mL; A=3 μg/mL. **(D)** Changes in the average MIC of the system over the simulations shown (A, B, and C).

## Discussion

The canonical distinction between bacteriostatic and bactericidal antibiotics has deeply influenced their clinical usage. Traditionally, bacteriostatic antibiotics were considered “weaker drugs”, but this traditional view is questionable. Drugs which are classified as bacteriostatic can and do kill bacteria in a concentration dependent manner ^22^. Moreover, meta-analysis studies do not demonstrate differences in the clinical success of therapy with either types of drugs even in severe infections ^11^. Indeed, bactericidal agents could be reserved for life-threatening infections, particularly in immunocompromised patients or those suffering from chronic infections. A more extended use of bacteriostatic drugs could be beneficial to spare the use and overuse of bactericidal antibiotics which fosters resistance. A limitation to progress in the extended use of bacteriostatic drugs is the shortage of pharmacodynamic (PD) data with these drugs.

The distinction between bacteriostatic and bactericidal antibiotics is confounded by another factor: The primary cellular and molecular targets do not necessarily differ between these classes of drugs. Several bacteriostatic antibiotics have mechanisms of action that one would anticipate being bactericidal: mecillinam and cefsulodin inhibit cell wall synthesis ^23^; novobiocin inhibits DNA gyrase ^24^; and rifampin inhibits the DNA-dependent RNA polymerase which is bacteriostatic for *E. coli* and bactericidal for *Mycobacterium* ^25,26^. Most notably, drugs which target the ribosome and inhibit protein synthesis can be either bactericidal (such as gentamicin or tobramycin) or bacteriostatic (such as the macrolides, phenicols, tetracyclines, oxazolidinones, and spectinomycin) ^27^. Interestingly, the potential bactericidal effect of one of the ribosome-targeting drugs, chloramphenicol, is prevented by the production of (p)ppGpp in the exposed cells ^28^. This suggests that the difference between a bacteriostatic drug and a bactericidal one is a property of the treated cell rather than the antibiotic ^29^. All the above examples illustrate the need for a better understanding of the PD, population biology, and evolutionary biology of treatment with bacteriostatic antibiotics.

In this study, we present evidence that bacteriostatic drugs of two different classes (the phenicols and macrolides) inhibit the growth of *E. coli* for extended periods, i.e. 30 days, and moreover maintain the culture in a kind of stationary phase where the density of viable bacterial cells is stable. Although, unlike stationary phase we found there to be an abundance of the limiting resource, implying that the cultures remain drug-limited even after a month. Most interestingly, despite the fact that the bacteria in the population appears to not be replicating due to the lack of net growth, evolution still occurred. A population of small colony variants (SCVs) emerged and ascended to become a significant population of bacteria. We attribute this evolution to the fact that even though the population at large was neither increasing nor decreasing in the cultures with bacteriostatic drugs, the population was replicating at a rate roughly equal to that at which it was dying. This finding is unanticipated and inconsistent with the common perception that bacteriostatic drugs simply arrest bacterial growth. This result questions the definition of bacteriostasis.

Our mathematical and computer-simulation model demonstrates that in the presence of an antibiotic, even when starting with an antibiotic susceptible population that generates resistant sub-populations at an extremely low rate, the resistant sub-population will ascend to dominate. When the selection pressure is removed this resistant sub-population will increase in susceptibility, however the population which then emerges to dominance is not the initial antibiotic susceptible population but is instead a population which readily generates resistant sub-populations. Stated another way, our model illustrates and our results confirm how HR can evolve in a population otherwise lacking this trait.

These SCVs were found to be highly resistant not only to the challenging agents, but even to some types of bactericidal agents, such as aminoglycosides and rifampin. Curiously, the resistance of the SCVs to chloramphenicol (CHL) and azithromycin (AZM) are not due to the canonical resistance mechanisms ^16,30^. SCVs have been implicated in treatment failure, primarily in *Staphylococcus aureus*, but there are limited reports of SCVs being associated with treatment failure in *E. coli* ^31,32^. Although SCVs of *E. coli* have been observed for a long time, they are under described both in their genetic basis and their role in bacterial communities. Two clinical isolates of *E. coli* SCVs have been studied, both of which have distinct metabolic disruptions by mutations in heme and thymidine biosynthesis pathways ^31,32^. Lab-generated SCVs have been described with mutations in Q8, hemin, and lipoic acid biosynthesis ^33-35^. However, none of these pathways appear to be disrupted in our SCVs.

We have yet to determine the genetic and molecular basis of the observed SCVs, but they appear to be distinct from the previously described mechanisms ^33-37^. Certain mutations observed here (Supplementary Table 1) might account for changes in MICs in the SCVs. For instance, in the case of AZM-induced SCVs, missense variants of the *citG* gene, encoding the 2-(5’’-triphosphoribosyl)-3’-dephosphocoenzyme-A synthase, were consistently found. This mutation might alter members of the GntR family of transcriptional regulators which could influence the DNA-binding properties of the regulator resulting in repression or activation of transcription, or could directly impact ATP synthesis ^38^, leading to the generation of the SCVs. We did find extra mutations in CHL C, AZM A, and AZM C involving the AcrAB efflux pump, 50S ribosomal proteins L4 and L22, and the Lon protease, all of which can modify the antibiotic susceptibility. The mutations in AZM C found in acrAB could interfere with AZM membrane transport ^39^. The mutations found in CHL C and AZM A in the gene encoding the 50S ribosomal protein L4 (a known chloramphenicol binding site) and L22, respectively, are a known mechanism of macrolide resistance ^40,41^. The mutation in the Lon protease in AZM A is frequently associated with changes in susceptibility as well ^42^. Taken together, these changes could account for the increase in the MIC to AZM and CHL but cannot account for the emergence of the SCVs. We found a mutation in a CHL-induced SCV in the gene encoding the Tyrosine-tRNA ligase. This mutation could account for the generation of SCVs, since inhibitors of this ligase strongly decrease bacterial growth, but this mutation was only found in one of the six SCVs ^43^. Most interestingly, we were unable to find any SNPs in one of the isolated CHL SCVs. However, all mutations found in the antibiotic-resistant SCVs were also found the in the antibiotic-suspectable revertants (Supplementary Table 2), thus even if the mutations presented could account for the transition to resistance, they cannot account for the transition to susceptibility. This is also true of the transition to and from the SCV phenotype. We cannot exclude that the transition from normal cells to SCVs could be due to a kind of “structural epistasis” (not a genetic one) due to the altered interactions between intracellular machinery and molecules when the shape of the cell is altered by antibiotics ^44^.

Unexpectedly, the antibiotic resistance observed here is transient, as would be anticipated for heteroresistance (HR), suggesting a high fitness cost of the mutations detected. In support of this HR hypothesis we found that these SCVs meet all the criteria set forth for HR: there are subpopulations present at a frequency greater than 10^−7^, with an MIC higher than 8x that of the main population, and reversion of the resistant subpopulation occurs in short order ^21^. To our knowledge, this is the first report of both the spontaneous evolution of HR as well as HR to bacteriostatic drugs.

## Methods

### Bacterial Strains

*E. coli* MG1655 was obtained from the Levin Lab bacterial collection ^13^. pAM34 with the origin of replication under control of an IPTG promoter and an ampicillin resistance cassette was obtained from Calin Guet from IST Austria ^18^.

### Growth Media

LB (Lysogeny Broth) (244620) was obtained from BD. The DM (Davis Minimal) minimal base without dextrose (15758-500G-F) was obtained from Sigma Aldrich (7 g/L dipotassium phosphate, 2 g/L monopotassium phosphate, 0.5 g/L sodium citrate, 0.1 g/L magnesium sulfate, 1 g/L ammonium sulfate). MHII plates were made from MHII broth (212322) obtained from BD. Glucose (41095-5000) was obtained from Acros. LB agar (244510) for plates was obtained from BD.

### Growth Conditions

Unless otherwise stated, all experiments were conducted at 37°C with shaking.

### Sampling bacterial densities

The densities of bacteria were estimated by serial dilution in 0.85% saline and plating. The total density of bacteria was estimated on LB agar plates.

### Antibiotics

Chloramphenicol (23660) was obtained from United States Pharmacopeia. Azithromycin (3771) was obtained from TOCRIS. Ampicillin (A9518-25G) was obtained from Sigma Aldrich. Isopropyl β-D-1-thiogalactopyranoside (IPTG; I56000-5.0) was obtained from Research Products International. All E-tests were obtained from bioMérieux.

### Estimating Minimal Inhibitory Concentrations

Antibiotic MICs were estimated using E-tests on MHII plates or via broth microdilution ^45,46^.

### Long Term Experiments

Flasks containing 10 mL of DM with 1000 μg/mL of glucose and an initial density of 10^5^ CFU/mL cells were grown at 37ºC with shaking for 30 days without transferring. For each biological replicate, initial inoculums were obtained from independent overnights of *E. coli* cultures. *E. coli* MG1655 was grown with either no drug, 4x MIC of CHL or 3x MIC of AZM. Samples were taken every 5 days and plated on LB agar plates. Twenty SCVs from each condition were isolated from the last time point and grown in the absence of antibiotic. Only those whose MIC was higher than the ancestral MG1655 after two overnight passages in the absence of the drug were selected.

### Long Term Experiments with non-replicative plasmid

Flasks containing 10 mL of DM with 1000 μg/mL of glucose and an initial density of 10^5^ CFU/mL cells were grown at 37ºC with shaking for 30 days. *E. coli* MG1655 pAM34 was grown with either 4x MIC of CHL, 3x MIC of AZM, ampicillin, or no selection. All experiments were performed in the absence of IPTG. pAM34 is a single copy number plasmid. Control Samples were taken every 5 days and plated on LB agar plates as well as 100 μg/mL Ampicillin and 0.5 mM IPTG LB agar plates.

### Sequencing

Complete genomes were obtained with hybrid Illumina/Nanopore sequencing by SeqCenter. Samples were extracted from single colonies using Zymo Quick-DNA HMW MagBead Kit. Oxford Nanopore Sequencing library prep was performed with PCR-free ligation library prep using ONT’s V14 chemistry. Long read sequencing was performd using R10.4.1 flowcells on a GridION with basecalling performed by Guppy in Super High Accuracy mode. Illumina libraries were prepared and sequenced per SeqCenter’s standards. Quality control and adapter trimming was performed with bcl-convert and porechop for Illumina and ONT sequencing respectively. Hybrid assembly with Illumina and ONT reads was performed with Unicycler ^47^. Assembly statistics were recorded with QUAST ^48^. Assembly annotation was performed with Prokka ^49^.

### Beta-galactosidase assay

The BetaRedTM β-Galactosidase Assay Kit (Sigma Aldrich) was used to experimentally measure the translation level. Flasks containing 10 mL of DM with 1000 μg/mL of glucose and an initial density of 10^5^ CFU/mL cells were grown at 37 ºC with shaking overnight in the absence or presence of different concentrations of CHL or AZM. The manufacturer protocol was followed, and OD 570 nm was measured. The experiments were performed in triplicate.

### Growth Rate Estimation

Exponential growth rates were estimated from changes in optical density (OD600) in a Bioscreen C. For this, 24-hours stationary phase cultures were diluted in LB or glucose-limited liquid media to an initial density of approximately 105 cells per mL. Five replicas were made for each estimate by adding 300μl of the suspensions to the wells of the Bioscreen plates. The plates were incubated at 37°C and shaken continuously. Estimates of the OD (600nm) were made every five minutes for 24 hours in LB and 48 hours in glucose-limited medium. Normalization, replicate means and error, growth rate, lag and maximum OD were found using a novel R Bioscreen C analysis tool accessible at https://josheclf.shinyapps.io/bioscreen_app.

### Residual Growth

After 30 days the cultures were centrifuged and filtered through a 0.22 μm filter. Strain resistant for CHL (Strain 1012 from the US Center for Disease Control’s MuGSI Isolate Bank which is *cmlA5* positive), AZM (Strain 1007 from the US Center for Disease Control’s MuGSI Isolate Bank which is *mph(A)* positive), or just MG1655 were added to the supernatants and allowed to grow for 24h.

### Antibiotic stability

Initial (day 0) and final (day 30) cultures from the long term experiment in the presence of CHL or AZM were centrifugated and the supernatants filtrated. Twenty μL of these supernatants were spotted on 7 mm blank disk and MG1655 was used as the lawn. After 24h of incubation, the zone of inhibition was recorded.

### Population Analysis Profile tests

Single colonies of the corresponding strain were inoculated into 10 mL of LB and grown overnight at 37 °C with shaking. Cultures were serially diluted in saline and all dilutions (10^0^ to 10^−7^) plated on LB agar plates containing 0, 1, 2, 4, 8, and 16 times the MIC of CHL or AZM. Plates were grown at 37 °C for 48 hours before the density of surviving colonies was estimated.

### Numerical Solutions (Simulations)

For our numerical analysis of the coupled, ordered differential equations presented (Supplementary Equations 1-12) we used Berkeley Madonna with the parameters presented in the captions of Figure 7 and Supplementary Figures 8 and 9. Copies of the Berkeley Madonna programs used for these simulations are available at www.eclf.net. Parameters were obtained from ^21,50,51^. To calculate the average MIC in our simulations, we take a weighted average of the MIC of each population.

## Supporting information

All supplemental materials

## Data Availability

All the data generated are available in this manuscript and its supporting supplementary material. The genomic data generated in this study have been deposited in the NCBI database under accession code PRJNA1032893 (https://www.ncbi.nlm.nih.gov/bioproject/?term=PRJNA1032893). Source data are provided with this paper.

## Acknowledgments

We would like to thank Danielle Steed for her discussion of the clinical implications and relevance of this work. We would also like to thank Jason Chen and LM Bradley for their unwavering assistance and support. We would also like to thank the other members of the Levin Lab. BRL would like to thank the US National Institute of General Medical Sciences for their funding support via R35 GM 136407 and the US National Institute of Allergy and Infectious Diseases for their funding support via U19 AI 158080-02.

## Author Contributions Statement

Conceptualization: TGG, BAB, JAM, FB, BRL

Methodology: TGG, BAB, JAM, ICM, BRL

Investigation: TGG, BAB, JAM, APS, ICM

Visualization: TGG, JAM

Funding Acquisition: BRL

Project Administration: BRL

Supervision: BRL

Writing– Initial Draft: TGG, BAB, JAM, APS, FB, BRL

Writing– Review & Editing: TGG, BAB, JAM, APS, ICM, FB, BRL

