## Supplementary material for "The Evolution of Heteroresistance via Small Colony Variants in *Escherichia coli* Following Long Term Exposure to Bacteriostatic Antibiotics": All supplemental materials

**Supporting Information for**
**The Evolution of Heteroresistance and Small Colony Variants in *Escherichia coli* Following Long Term Bacteriostatic Drug Exposure**

Teresa Gil-Gil, Brandon A. Berryhill, Joshua A. Manuel, Andrew P. Smith, Ingrid C. McCall, Fernando Baquero, Bruce R. Levin*

*Corresponding Author


**SUPPLEMENTARY TEXT**
**Models for Non-replicating Plasmid**

To explain how the results with the non-replicating plasmid in Figure 2 are consistent with our hypothesis that bacteria confronted with bacteriostatic antibiotics are replicating despite the absence of net growth, we present two models below. The first model parallels the long term exposure to bacteriostatic drugs while the second model parallels the control experiment in the absence of any drug.

*Plasmid in the presence of a bacteriostatic drug*

In this model we assume the net growth rate of the population at large is zero. We also assume that the plasmid bearing cells are not capable of increasing in absolute number, but instead decrease in frequency by transitioning to cells not bearing the plasmid at half of the growth rate. For this model, the replicating plasmid- free cells are labeled N in cells per mL which grow at some rate v, per cell per hour. The cells containing the plasmid, P cells per mL, also grow at the same rate v, per cell per hour. Since half of the divisions of the P population result in loss of the plasmid due to segregation loss, the transition between states P and N is effectively half the growth rate. Equations 1 and 2 detail this relationship.

$$\frac{dN}{dt} = 0.5 \cdot v \cdot P \quad \textbf{Eq. 1}$$
$$\frac{dP}{dt} = -0.5 \cdot v \cdot P \quad \textbf{Eq. 2}$$

*Plasmid in the absence of a bacteriostatic drug*

In this model we assume that all cells can grow, but since half the cells per division of P lose the plasmid and become N, the absolutely number of P remains constant while the frequency of P decreases. We also assume that there is an absolute carrying capacity such that when the total density of cells reaches that capacity all transitions and growth stops. All other assumptions for this model are as above. This relationship is giving by Equations 3 and 4.

$$\frac{dN}{dt} = (N \cdot v + 0.5 \cdot v \cdot P) \quad \textbf{Eq. 3}$$
$$\frac{dP}{dt} = 0 \quad \textbf{Eq. 4}$$

*Model predictions*

In Supplemental Figure 3 we illustrate with numerical solutions to the predicted results from the above equations. Shown in both panels in black are plasmid-free cells, and shown in blue is the frequency of plasmid-bearing to total cells as given by (P/P+N).

### Evolution of Heteroresistance Model

In the absence of antibiotics, the maximum growth rates of these populations are, respectively, $v_{maxN}$, $v_{maxS}$, and $v_{maxH}$ (>0 per cell per hour). In the presence of antibiotics, the minimum growth rate (maximum death rates) of these populations are $v_{minN}$, $v_{minS}$, and $v_{minH}$ (<0 per cell per hour), and the respective MICs of these populations are $MIC_N$, $MIC_S$, and $MIC_H$ µg/mL. The net rates of growth of these three populations are proportional to the concentration of the resource, $r$ µg/mL and the concentration of the antibiotic, $A$ µg/mL [1,2]. The equations below are given in the general form where $X$ is either N, S, or H.

$$\Pi_X(A,r) = v_{maxX} - \left[\frac{(v_{maxX}-v_{minX})\cdot\left(\frac{A}{MIC_X}\right)^{Kx}}{\left(\frac{A}{MIC_X}\right)^{Kx}-\left(\frac{v_{minX}}{v_{maxX}}\right)}\right]\cdot \psi x(r) \quad \textbf{Eq.5}$$

$$yx(r) = \frac{r}{(r+k_x)} \quad \textbf{Eq.6}$$

$K_x$, the Hill coefficient [1], is a shape parameter where the greater the value of $K_x$ the more acute the function. The parameter $k_x$, the Monod constant, is the concentration of the resource when the growth rate is half its maximum value [3].

With the above definitions and assumptions, the rates of change in the densities of the populations of bacteria and the change in resource and antibiotic concentrations are given by the below set of coupled differential equations (Eq. 7-11).

$$\frac{dN}{dt} = \Pi_N(A,r)\cdot(N + N\cdot(\mu_{sn} - \mu_{ns})) \quad \textbf{Eq. 7}$$

$$\frac{dS}{dt} = \Pi_S(A,r)\cdot(S + S\cdot(\mu_{ns} - \mu_{sn} + \mu_{hs} - \mu_{sh})) \quad \textbf{Eq. 8}$$

$$\frac{dH}{dt} = \Pi_H(A,r)\cdot(H + H\cdot(\mu_{sh} - \mu_{hs})) \quad \textbf{Eq. 9}$$

$$\frac{dr}{dt} = -e\cdot(N\cdot v_{maxN} + S\cdot v_{maxS} + H\cdot v_{maxH}) \quad \textbf{Eq. 10}$$

$$\frac{dA}{dt} = -da\cdot A \quad \textbf{Eq. 11}$$

The conversion efficiency, $e$ µg [4], is the amount of the limiting resource needed to produce a new cell (Eq. 10) and the parameter $da$ is the hourly rate of decline in the concentration of the antibiotic in µg/hour (Eq. 11).

We use Berkeley Madonna and the Euler method to generate numerical solutions to these differential equations. In these simulations the changes in the densities of these populations and concentrations of the limiting resource are deterministic. The generation of mutants, however, is stochastic and simulated by a Monte Carlo process [5]. At each time interval, t to dt (where dt is the step size), a random number (0≤ z ≤1) from a rectangular distribution is generated. If the random number is less than the probability in Equation 12, we add 1/dt to the $X$ population. For Equation 12, Vol is the volume of the vessel which we simulate as 10 mL and $X$ is the density of the respective population.

$$P(\mu) = \mu_X \cdot X \cdot dt \cdot \mathrm{Vol} \quad \textbf{Eq. 12}$$

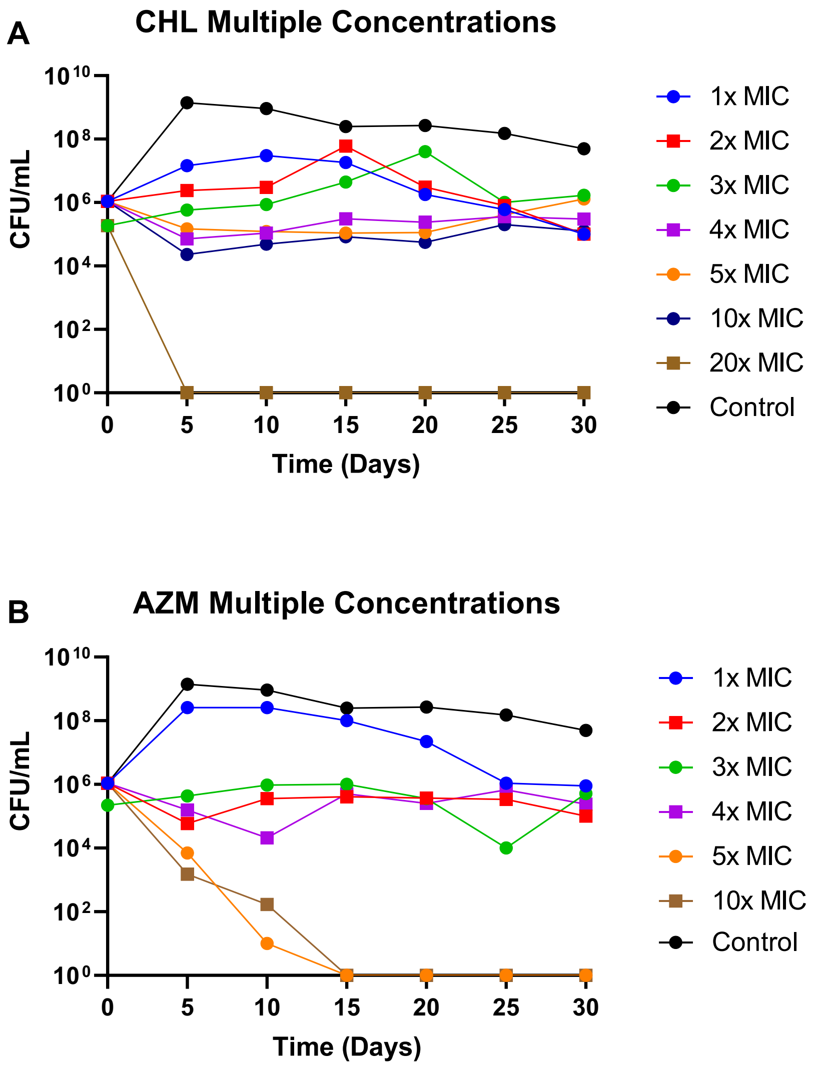


**Supplementary Figure 1. Selection of bacteriostatic concentrations of CHL and AZM.** CFU/mL of *E. coli* MG1655 in cultures with varying concentrations of **(A)** CHL or **(B)** AZM for 30 days. Blue line- 1x MIC, Red line- 2x MIC, Green line- 3x MIC, Purple line- 4x MIC, Orange line- 5x MIC, Brown line- 10x MIC, Black line- Drug-free control.

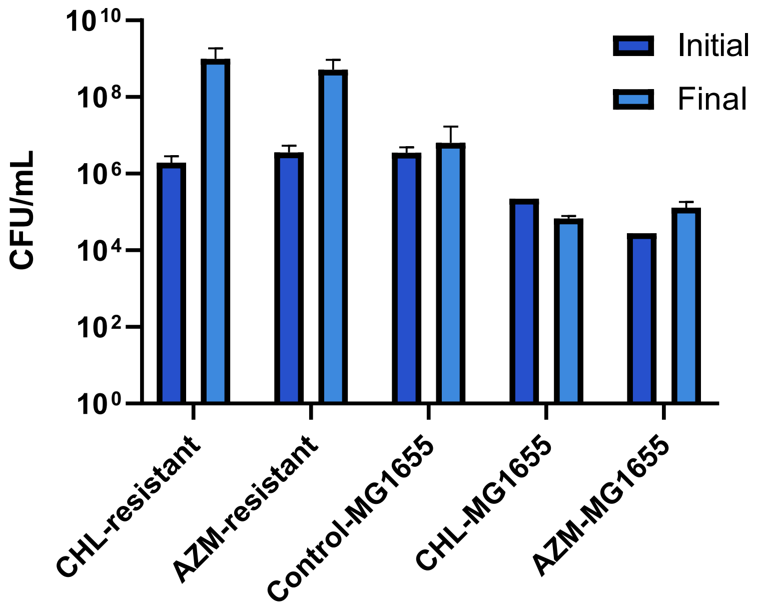


**Supplementary Figure 2. Growth of *E. coli* strains in supernatants from day 30 of the long term experiments**. 5x$10^6$ CFU/mL of AZM-resistant, CHL-resistant, or *E. coli* MG1655 strains when inoculated in the cell-free supernatants of the 30-day time point. The cell-free supernatants were plated on LB agar plates and no colonies were found. *E. coli* MG1655 was inoculated on cell-free supernatants from the control and both antibiotics and AZM-resistant or CHL-resistant strain on cell-free supernatants in which the corresponding antibiotic was present. Error bars represent the standard deviation of 4 biological replicates. Dark blue- Initial (Time= 0 hours), Light blue- Final (Time= 24 hours).

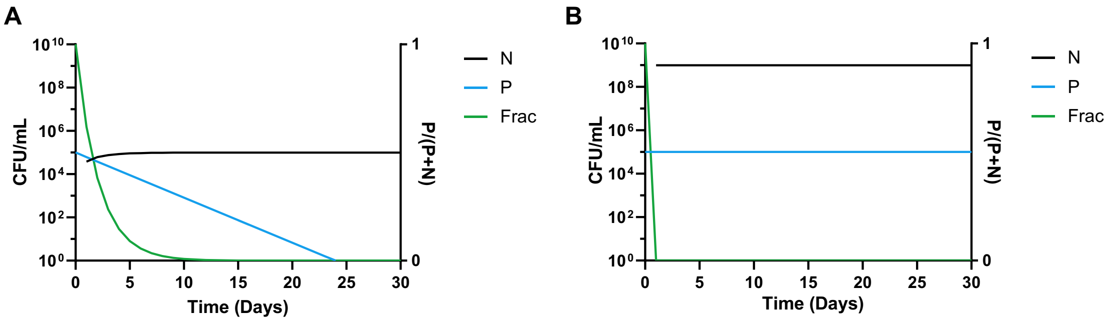


**Supplementary Figure 3. Numerical Solutions for Simulations of The Non-replicating Plasmid.** In both panels, Initial N= 0 cells per mL, Initial P=$10^5$ cells per mL, v= 0.04 per cell per hour (or one division per cell per day as calculated from Figure 2), and we assume a maximum density of $10^9$ cells per mL. Shown on the left axis is CFU/mL and on the right axis "Frac" which is P/(P+N). **A** Simulations in the presence of bacteriostatic drug. **B** Simulations in the absence of bacteriostatic drug.

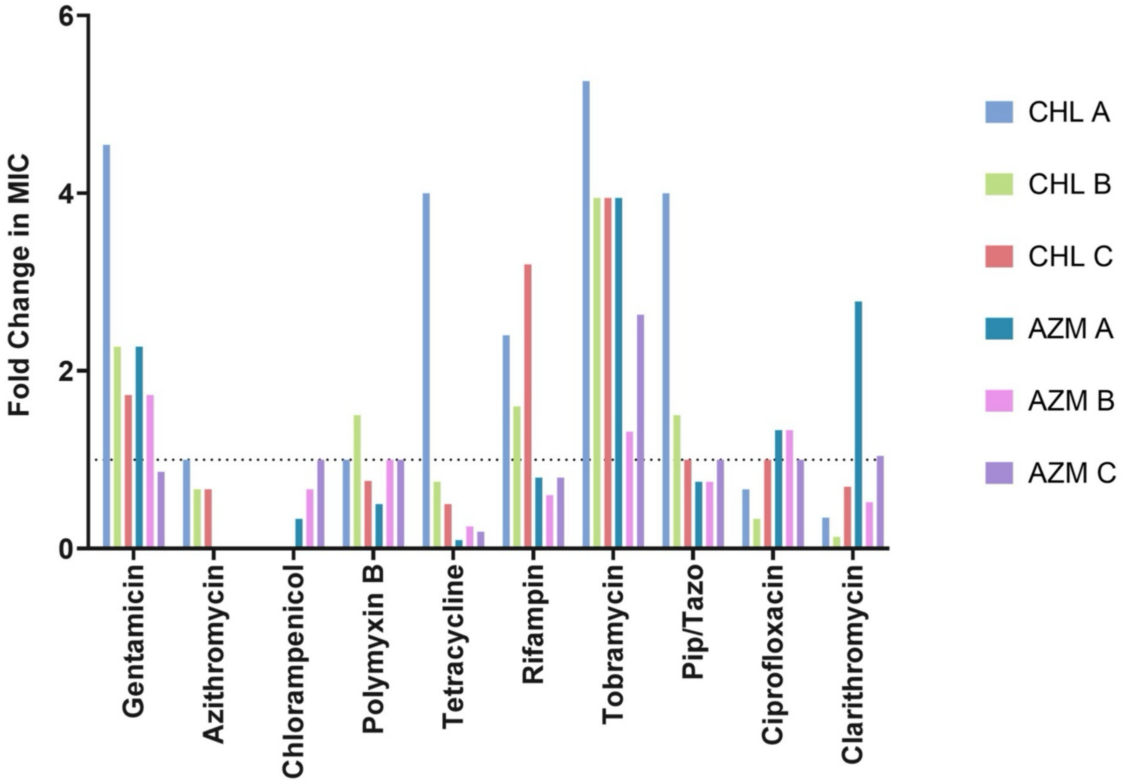


**Supplementary Figure 4. SCV collateral susceptibilities and cross-resistances.** Ratio of the MICs of the 6 SCVs compared to the ancestral *E. coli* MG1655 strain measured by E-test. Pip/Tazo stands for piperacillin/tazobactam.

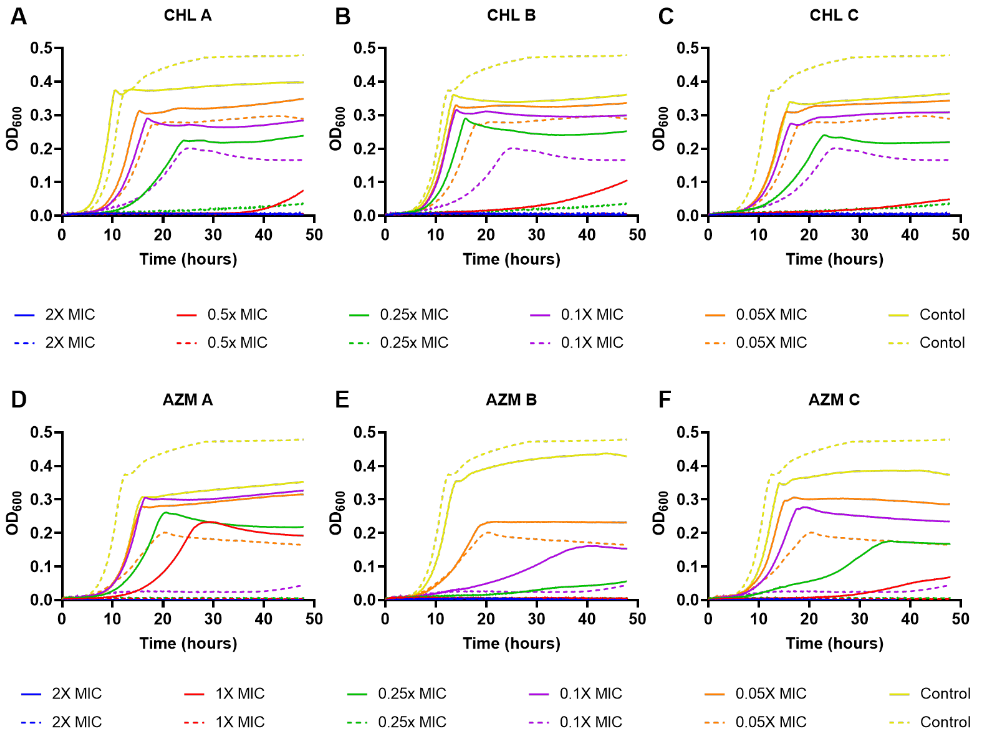


**Supplementary Figure 5. Growth dynamics of CHL SCVs (solid lines A-C), AZM SCVs (solid lines D-F) and *E. coli* MG1655 (dashed lines A-F).** Changes in optical density (600nm) of *E. coli* MG1655 exposed to five different concentrations of both drugs for 48 hours in minimal media. Lines are representative of the average of five technical replicas and normalized to the time zero optical density. Each concentration is shown as a fraction of the MIC of the SCV for the noted drug.

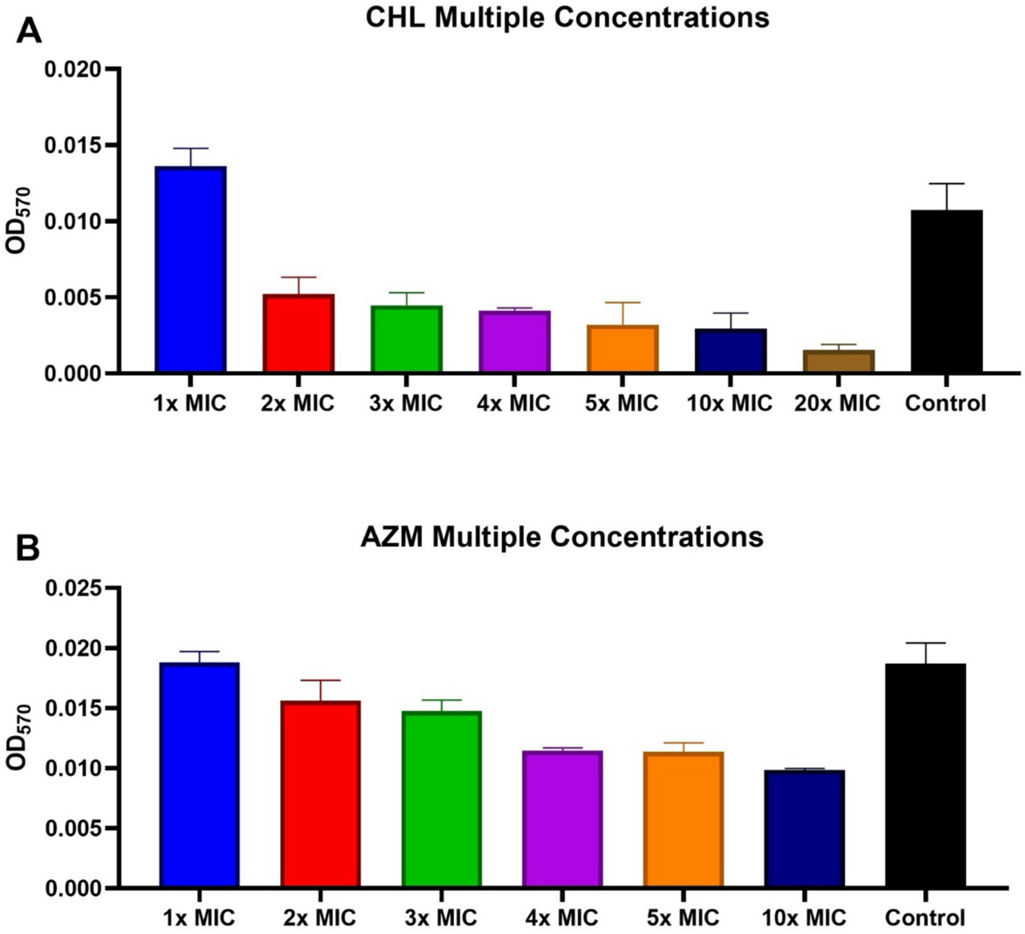


**Supplementary Figure 6. An assay of beta-galactosidase as a proxy for translation level during antibiotic exposure.** The beta-galactosidase levels as determined by absorbance for the ancestral *E. coli* MG1655 at different concentrations of (A) CHL and (B) AZM. Shown are means and standard deviations of three independent, biological replicates.

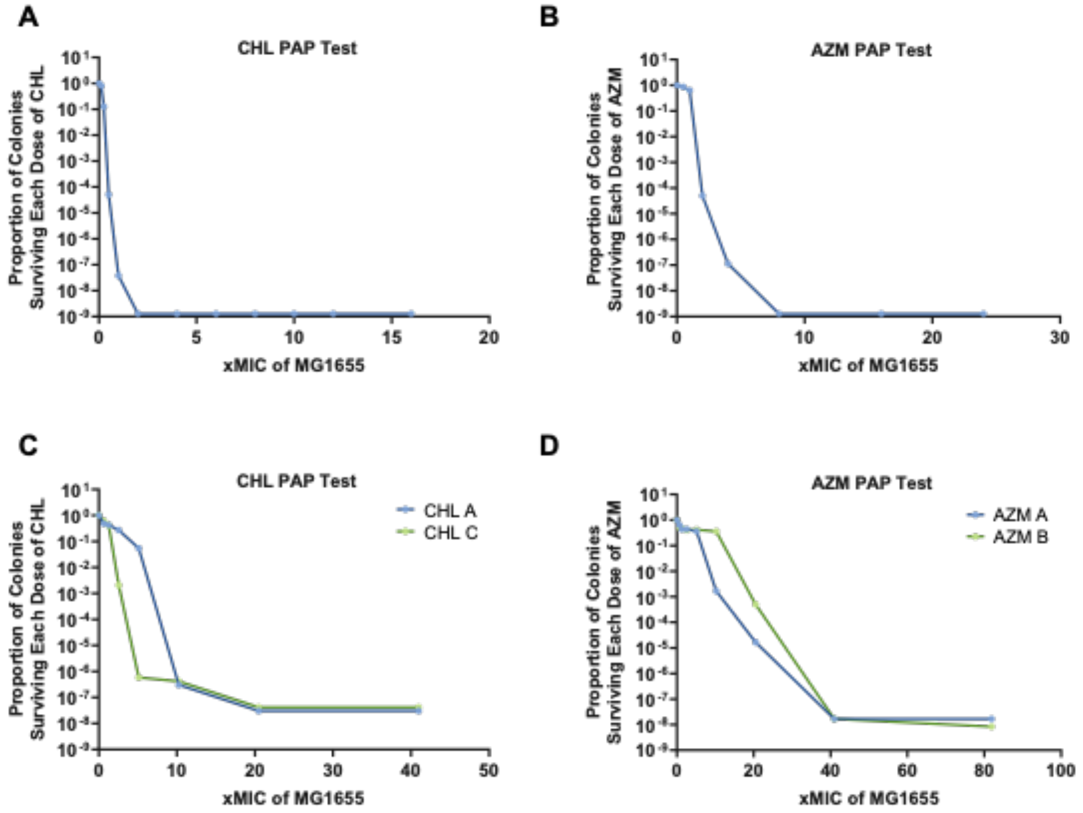


**Supplementary Figure 7. PAP tests. (A)** PAP test of *E. coli* MG1655 with CHL. **(B)** PAP test of *E. coli* MG1655 with AZM. **(C)** PAP test of CHL SCVs. **(D)** PAP test of AZM SCVs.

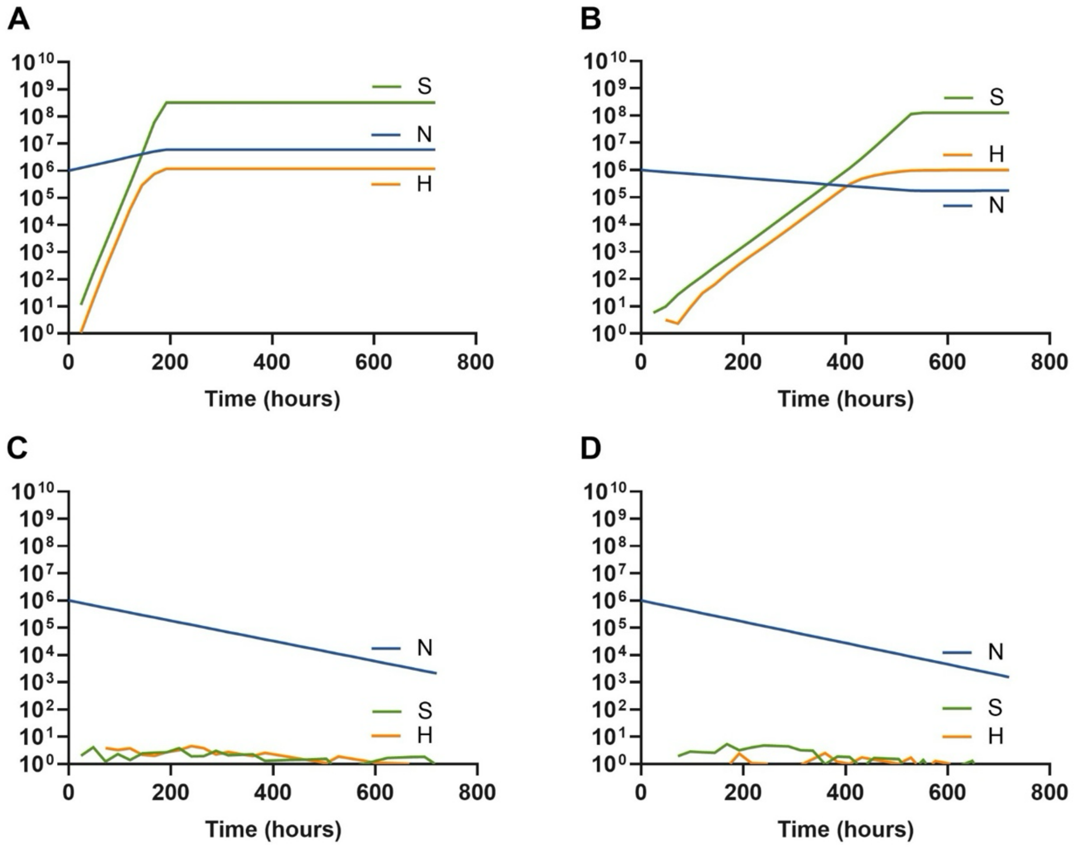


**Supplementary Figure 8. Computer simulations of the proposed model of evolved HR with differing antibiotic concentrations.** Parameters used for these simulations are $e$= 5x10$^{-7}$ µg/cell; $v_{maxN}$ = 1.0, $v_{maxS}$ = 0.5, $v_{maxH}$ = 1.0 per cell per hour; $v_{min}$ = -0.01 per cell per hour; K=1; k=1; $MIC_N$=1.0, $MIC_S$=8.0, $MIC_H$=2.0 µg/ml; $da$= 0 µg/hour; $\mu_{ns}$=1x10$^{-8}$, $\mu_{sn}$=1x10$^{-8}$, $\mu_{sh}$=1x10$^{-3}$, $\mu_{hs}$= 1x10$^{-3}$ per cell per hour. **(A)** A = 0.5 µg/mL. **(B)** A = 1.5 µg/mL. **(C)** A =7 µg/mL, note the S population ascends at 1400 hours. **(D)** A= 10 µg/mL, note the S population fails to ascend even at 1400 hours.

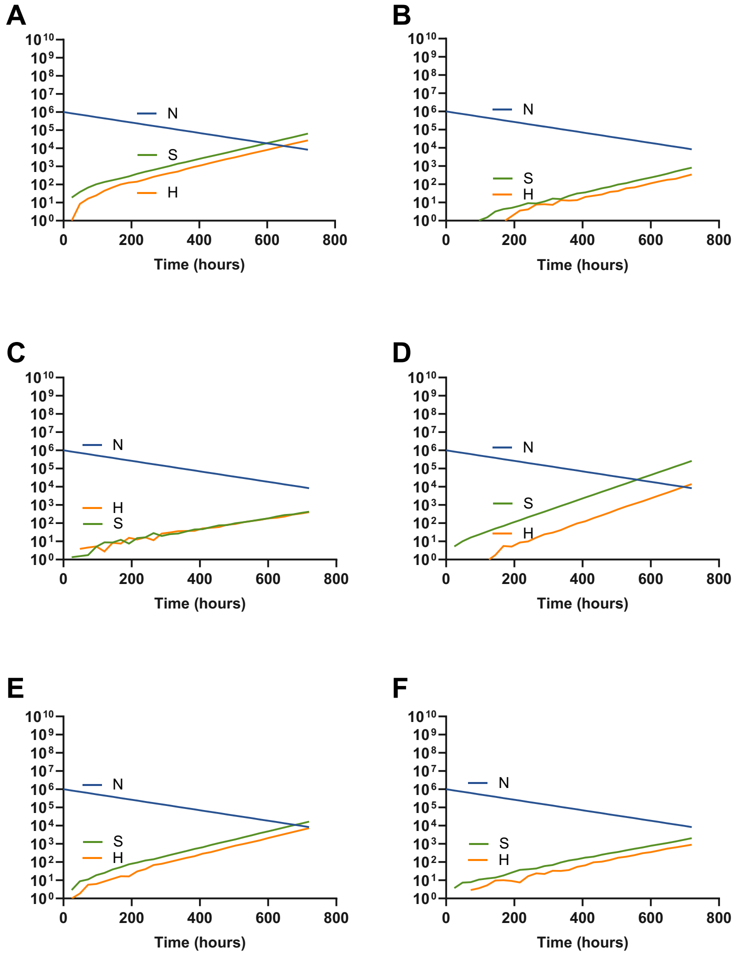


**Supplementary Figure 9. Computer simulations of the proposed model of evolved HR with differing transitions rates and fitness costs.** H and S selection from an initial sensitive population. Common parameters used for these simulations are $e$= 5x10$^{-7}$ µg/cell; $v_{maxN}$ = 1.0, $v_{maxH}$ = 1.0 per cell per hour; $v_{min}$ = -0.01 per cell per hour; K=1; k=1; $MIC_N$=1.0, $MIC_S$=8.0, $MIC_H$=2.0 µg/ml; $da$= 0 µg/hour; **(A)** Parameters used for these simulations are $v_{maxS}$ = 0.5 per cell per hour; $\mu_{ns}$=1x10$^{-7}$, $\mu_{sn}$=1x10$^{-7}$, $\mu_{sh}$=1x10$^{-3}$, $\mu_{hs}$= 1x10$^{-3}$ per cell per hour. **(B)** Parameters used for these simulations are $v_{maxS}$ = 0.5 per cell per hour; $\mu_{ns}$=1x10$^{-9}$, $\mu_{sn}$=1x10$^{-9}$, $\mu_{sh}$=1x10$^{-3}$, $\mu_{hs}$= 1x10$^{-3}$ per cell per hour. **(C)** Parameters used for these simulations are $v_{maxS}$ = 0.5 per cell per hour; $\mu_{ns}$=1x10$^{-8}$, $\mu_{sn}$=1x10$^{-8}$, $\mu_{sh}$=1x10$^{-2}$, $\mu_{hs}$= 1x10$^{-2}$ per cell per hour. **(D)** Parameters used for these simulations are $v_{maxS}$ = 0.5 per cell per hour; $\mu_{ns}$=1x10$^{-8}$, $\mu_{sn}$=1x10$^{-8}$, $\mu_{sh}$=1x10$^{-4}$, $\mu_{hs}$= 1x10$^{-4}$ per cell per hour. **(E)** Parameters used for these simulations are $v_{maxS}$ = 1 per cell per hour; $\mu_{ns}$=1x10$^{-8}$, $\mu_{sn}$=1x10$^{-8}$, $\mu_{sh}$=1x10$^{-3}$, $\mu_{hs}$= 1x10$^{-3}$ per cell per hour. **(F)** Parameters used for these simulations are $v_{maxS}$ = 0.1 per cell per hour; $\mu_{ns}$=1x10$^{-8}$, $\mu_{sn}$=1x10$^{-8}$, $\mu_{sh}$=1x10$^{-3}$, $\mu_{hs}$= 1x10$^{-3}$ per cell per hour.

**Supplementary Table 1. SCV Genomic Changes**

| SCV | GENE | NUCLEOTIDE POSITION | TYPE | NUCLEOTIDE CHANGE | EFFECT | PRODUCT |
|---|---|---|---|---|---|---|
| **CHL B** | *tyrS* | 2166796 | Ins | T → TTAACGG | Conservative in-frame insertion<br><br>Asn387→Gly388dup | Tyrosine-tRNA ligase |
| **CHL C** | *rplD* | 431632 | SNP | A → G | Missense variant<br><br>Lys63Arg | 50S ribosomal protein L4 |
| | | 3178132 | SNP | C → A | | Unannotated region |
| **AZM A** | *rplV* | 433741 | Del | ATGAAGCGCA TTATGCCGCGT GCAAAAGGTC GTGCAGATCG CATCC → A | Disruptive in-frame deletion<br><br>Ile85Arg99del | 50S ribosomal protein L22 |
| | *lon_1* | 2864809 | SNP | C → A | Stop gained<br><br>Ser422 | Lon protease |
| **AZM B** | *citG* | 3235035 | SNP | A → G | Missense variant<br><br>Glu234Gly | 2-(5''-triphosphoribosyl)-3'-dephosphocoenzyme-A synthase |
| | *citG* | 3235041 | SNP | G → T | Missense variant<br><br>Gly236Val | 2-(5''-triphosphoribosyl)-3'-dephosphocoenzyme-A synthase |
| **AZM C** | *citG* | 3235035 | SNP | A → G | Missense variant<br><br>Glu234Gly | 2-(5''-triphosphoribosyl)-3'-dephosphocoenzyme-A synthase |
| | *citG* | 3235041 | SNP | G → T | Missense variant | 2-(5''-triphosphoribosyl)-3'-dephosphocoenzyme-A synthase |

| | | | | | | |
|---|---|---|---|---|---|---|
| | | | | | Gly236Val | |
| | *acrB_2* | 3399923 | SNP | G → T | Missense variant<br><br>Gly236Val | Multidrug efflux pump subunit AcrB |

Ins: insertion; SNP: Single Nucleotide Polymorphism; Del: deletion

**Supplementary Table 2. Genomic Changes of Revertants**

| SCV | GENE | NUCLEOTIDE POSITION | TYPE | NUCLEOTIDE CHANGE | EFFECT | PRODUCT |
|---|---|---|---|---|---|---|
| **CHL B** | *tyrS* | 2166796 | Ins | T → TTAACGG | Conservative in-frame insertion<br><br>Asn387→Gly388dup | Tyrosine-tRNA ligase |
| | ***rpoC*** | 4335806 | SNP | C → A | Missense variant<br><br>Asp410Tyr | DNA-directed RNA polymerase subunit beta |
| **CHL C** | *rplD* | 431632 | SNP | A → G | Missense variant<br><br>Lys63Arg | 50S ribosomal protein L4 |
| | | 3178132 | SNP | C → A | | Unannotated region |
| **AZM A** | *rplV* | 433741 | Del | ATGAAGCGCA TTATGCCGCGT GCAAAAGGTC GTGCAGATCG CATCC → A | Disruptive in-frame deletion<br><br>Ile85Arg99del | 50S ribosomal protein L22 |
| | *lon_1* | 2864809 | SNP | C → A | Stop gained<br><br>Ser422 | Lon protease |
| **AZM B** | ***rpoA*** | 443272 | SNP | C → A | Missense variant<br><br>Arg191Ser | DNA-directed RNA polymerase subunit alpha |
| | *citG* | 3235035 | SNP | A → G | Missense variant<br><br>Glu234Gly | 2-(5''-triphosphoribosyl)-3'-dephosphocoenzyme-A synthase |
| | *citG* | 3235041 | SNP | G → T | Missense variant | 2-(5''-triphosphoribosyl)-3'-dephosphocoenzyme-A synthase |

| | | | | | | |
|---|---|---|---|---|---|---|
| | | | | | Gly236Val | |
| | ***acrB_2*** | 3399923 | SNP | T → C | Missense variant<br><br>Leu828Ser | Multidrug efflux pump subunit AcrB |
| **AZMC** | *citG* | 3235035 | SNP | A → G | Missense variant<br><br>Glu234Gly | 2-(5''-triphosphoribosyl)-3'-dephosphocoenzyme-A synthase |
| | *citG* | 3235041 | SNP | G → T | Missense variant<br><br>Gly236Val | 2-(5''-triphosphoribosyl)-3'-dephosphocoenzyme-A synthase |
| | *acrB_2* | 3399923 | SNP | T → C | Missense variant<br><br>Leu828Ser | Multidrug efflux pump subunit AcrB |

Ins: insertion; SNP: Single Nucleotide Polymorphism; Del: deletion
